# Signals of recent tropical radiations in Cunoniaceae, an iconic family for understanding Southern Hemisphere biogeography

**DOI:** 10.1101/2020.01.23.916817

**Authors:** Ricardo A. Segovia, Andy R. Griffiths, Diego Arenas, A. A. Piyali Dias, Kyle G. Dexter

## Abstract

Extratropical angiosperm diversity is thought to have arisen from lineages that originated in the more diverse tropics, but studies of dispersal between tropical and extratropical environments and their consequences for diversification are rare. In order to understand the evolutionary effects of shifts between the tropics and extratropics, defined here as areas that *do* versus *do not* regularly experience freezing temperatures, we studied the biogeographic history and associated diversification patterns of Cunoniaceae. We mapped the distribution of all species in the family and combined this with a newly constructed phylogeny for the family. The family shows a long evolutionary association with both tropical and extratropical environments, the tropics house considerably greater species richness of Cunoniaceae. Indeed, both tropical and extratropical environments appear to have had a similar number of lineages until 12 Ma, after which time the number of lineages in tropical areas increased at a faster rate. In addition, community phylogenetic approaches show that tropical regions have markedly less phylogenetic diversity than expected given their species richness, which is further suggestive of recent species radiations in tropical areas. The Cunoniaceae show an atypical pattern for angiosperms of frequent shifts between tropical and extratropical environments, but despite this, shows a more conventional pattern of higher, although recent, diversification rates in the tropics. Our results support the idea that high angiosperm species richness in the tropics may result from the tropics acting as a cradle of recent angiosperm diversification.

## Introduction

The widely observed pattern of decreasing plant diversity from low to high latitudes is accompanied by conspicuous phylogenetic, taxonomic and functional turnover (*e. g.* [1, 2]). Within angiosperms, the evolutionary dispersal of tropical, low latitude lineages into extratropical, high latitudes is relatively rare [3, 4]. When such dispersal does occur, it is enabled by the evolution of functional traits conveying tolerance of freezing environments, such as herbaceous habit, deciduousness and hydraulic adjustments [5]. Similarly, the movement of temperate-adapted lineages back into truly tropical climates also seems rare [6]. As such, the modern distribution of angiosperm tree lineages across the Americas is strongly associated with the presence or absence of freezing temperatures [7]. Although the origin of extratropical plant diversity may be associated with the evolution of tolerance to freezing conditions [8], the persistent restriction of tropical lineages to tropical environments and extratropical lineages to extratropical environments [9] limits opportunities to study the evolutionary processes associated with shifts between tropical and extratropical environments (*e. g.* [10]). The angiosperm family Cunoniaceae offers one such rare opportunity to study the evolutionary history of a clade in the context of tropical-extratropical environmental shifts.

Cunoniaceae (Oxalidales) is an iconic Southern Hemisphere family, containing 28 genera and *c.* 300 species of trees and shrubs, inhabiting both tropical and extratropical environments [11]. The cross-continental distribution of genera such as *Cunonia*, *Weinmannia*, *Eucryphia* or *Caldcluvia* reflects a deep history in Southern Hemisphere landmasses. Indeed, the major diversification of the Cunoniaceae is thought to have occurred in the Gondwanan region [12]. However, whether tropical (*i. e.* frost-free environments) and/or extratropical (*i. e.* freezing environments) environments are associated with the initial and later diversification is currently unclear. Similarly, a tropical or extratropical origin and diversification of *Weinmannia*, the most speciose genus in the family (comprising ~44% of species), has yet to be clarified [13].

Biogeographic studies on the distribution of the Southern Hemisphere biota have traditionally centered on two main processes: vicariance associated with the Gondwanan breakup [14, 15, 16, 17, 18], and long-distance dispersal through wind and ocean currents [19, 20, 21]. Yet, recently documented evidence for environmental constraints on the evolution and dispersal of lineages [22, 23], underlines the need to develop an alternative perspective in the study of biogeography, based on the principle of phylogenetic niche or biome conservatism. Moreover, Southern Hemisphere trends suggest that the environmental divide between the tropics and extratropics may be a key determinant of the distribution of modern-day diversity. For example, phylogenetic analyses for lineages in the Southern Hemisphere show that intercontinental dispersal events are more frequent than environmental shifts, even between neighboring tropical and extratropical biomes [24]. Furthermore, the fossil record shows a lower than expected interchange of plant lineages between the Neotropics and the extratropics of southern South America across the whole Cenozoic [25].

Hypotheses regarding evolutionary biogeography can be directly explored using phylogenetic approaches. For example, ancestral state reconstructions can track the evolutionary history of lineage characteristics, such as climatic niches, across a phylogenetic tree [26, 27]. More recently, biogeographic insights have been provided by analyzing variation in community phylogenetic structure across biogeographic regions, biomes and environmental gradients (*e. g.* [28, 29, 30, 31]). Standardized phylogenetic diversity (sPD), that is phylogenetic diversity (PD) given variation in species richness (SR), can be used as a metric of community phylogenetic structure [32]. Areas with significantly lower PD than expected, given SR, are composed of closely related species. Such phylogenetic clustering of related species in an assemblage can result from rapid speciation, selective extinction of older lineages, or reduced interchange of lineages among communities or geographic regions [33, 34]. In contrast, areas with significantly greater PD than expected, given SR, are composed of distantly related species. Such phylogenetic overdispersion can result from lower diversification rates in recent time than deeper time, higher recent extinction, or frequent interchange of distantly related lineages among regions [33, 34].

In this study, we combine a newly derived phylogeny for Cunoniaceae with global occurrence and climatic data to gain insight into how dispersal events across the tropical-extratropical divide shaped the evolutionary biogeography of the family. We firstly reconstruct the ancestral tropical or extratropical affiliation of all 28 described genera within Cunoniaceae. In addition, we make a comparison of phylogenetic structure, phylogenetic signal, and variation in lineage diversity through time between tropical, non-freezing environments and extratropical, freezing environments. The evolutionary origins of angiosperms in general is thought to have occurred in tropical, non-freezing environments, with only a subset of lineages evolving tolerance to freezing conditions and making the shift to extratropical environments [8]. If the Cunoniaceae follow this general tropical-origin trend, we anticipate a phylogenetically clustered pattern in assemblages from extratropical environments and/or a phylogenetic signal for extratropicality (*i. e.* extratropical species will be more closely related than expected by chance). Alternatively, if tropical distribution is an evolutionary novelty within the Cunoniaceae, with tropical lineages deriving from extratropical lineages, we expect phylogenetic clustering in assemblages from tropical environments, and/or a phylogenetic signal for tropicality. Finally, we evaluate whether there is phylogenetic structure behind the variation in species richness of Cunoniaceae across climatic space. In environments where Cunoniaceae has experienced a recent and rapid species radiation, we expect assemblages to have both high species richness and display phylogenetic clustering (*i. e.* significantly low sPD values). In contrast, in environments where the diversification of Cunoniaceae has been comparatively slow, we expect random sPD values centred on zero, or a pattern of phylogenetic overdispersion (*i. e.* significantly high sPD values), if there are many old, yet undiversified lineages.

## Material and Methods

### Distribution dataset

Cunoniaceae occurrence data were downloaded from GBIF on August 28th, 2019 using the ‘occ_search’ function in the ‘rgbif’ package [35] for R [36]. First, we excluded duplicate and incomplete occurrences. Then, we used the function ‘clean_coordinates’ in the R package ‘CoordinateCleaner’ [37] to filter the occurrence data, excluding collections from capitals, centroids of countries, GBIF headquarters, natural history institutions, and collections occurring in the sea. We also identified records considered as outliers in the environmental space the given species occupies. Environmental outlier detection was performed by running a reverse-jackknife method on six climatic variables: mean annual temperature, maximum temperature of the warmest month, minimum temperature of the coldest month, mean annual precipitation, precipitation of the wettest month, and precipitation of the driest month, extracted from WorldClim [38]. These variables were selected to reflect the climatic averages and boundaries of species’ ranges. The method, which has precedence (*e. g.* [39, 40], consists of identifying outlier samples (*i. e.* occurrences) for each climatic variable based on a critical threshold derived from the mean, standard deviation, and range of the whole set of samples for a given species (*i. e.* all occurrence points). Potential outlier occurrences were determined as those where values for at least 2 of the 6 (> 20%) environmental variables were identified as statistical outliers. If there was not enough information to compute this analysis (here, less than 10 occurrences for a given species), then the environmental outlier analysis was not performed. The final GBIF distribution dataset contained 54,717 records, with an average of 181 records per species, a minimum of 1 record (for 19 spp) and a maximum of 26,432 records (for *Weinmannia racemosa*).

### Ecological Classification

Given the climatic [41] and evolutionary [7] importance of freezing environments as a division between tropics and extratropics, we classified the species in our dataset as Tropical, Extratropical or Generalist based on the proportion of occurrences subject to regularly freezing temperatures. We defined the tropics as those environments not subject to freezing temperatures in a normal year. The extratropics were conversely defined as those environments subjected to regular freezing temperatures. Using a global climatic layer for cumulative frost days over 117 years between 1901 and 2018 (Climatic Research Unit (CRU), [42]), we defined 117 total days of freezing (*i. e.* one day of frost per year on average) as the threshold for ‘regularly experiencing’ freezing temperatures versus not. Some regions can sum 117 days with freezing temperatures in a non-regular way, such as areas subject to periodic polar outbreaks [43]. However, we consider our approach to provide an objective threshold for environments where freezing is relevant. Based on these freezing data we defined Extratropical species as those with more than 75% of their occurrences in regions with regular freezing temperatures and Tropical species as those with less than 25% of occurrences in regions with regular freezing. Those species with a percentage between 25% and 75% of occurrences in regions with regular freezing were classified as “Generalists” (Fig. 1).

**Figure 1.**
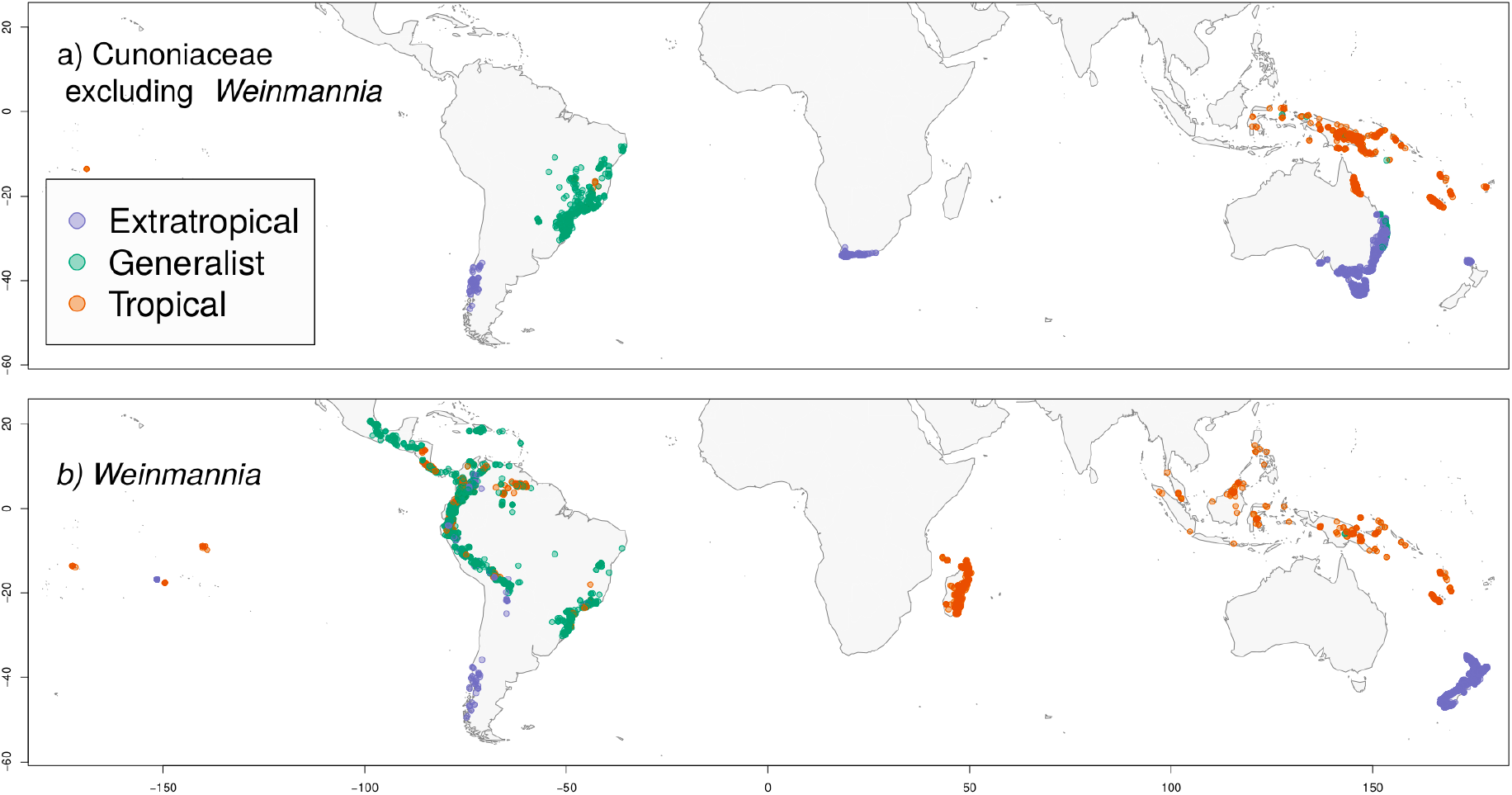
Distribution of the family Cunoniaceae. a) Distribution of the family Cunoniaceae excluding *Weinmannia* (169 non-Weinmannia spp) and b) Distribution of *Weinmannia* (133 spp). Purple dots represent occurrences of species classified as Extratropical. Orange dots represent occurrences of species classified as Tropical. Green dots represent occurrences of species classified as Generalist.

### Phylogeny Construction

In order to build the phylogeny, sequence data from NCBI were gathered using PyPHLAWD, with a clustering method [44]. The 6 best clusters (trnL (52 spp), phyC (45 spp), rbcL (42 spp), ncpGS (38 spp), GapC (20 spp), matK (15 spp)) were aligned individually in MAFFT [45]. The total matrix included 82 species from 22 genera, and 7,179 bps. A Maximum Likelihood tree was estimated using RaxML v8.2.12 [46], with a separate GTR + Γ model for each gene. This resulting phylogeny was then temporally calibrated using treePL [47], with the crown age for the family taken from the its first occurrence in the fossil record: Maastrichtian [48]. Thus, we set crown age to a maximum of 70 Myr and a minimum of 66 Myr.

We manually added genera to the phylogeny for which no DNA sequences were available. Exact placement of manual additions was based on consultation of morphological and molecular phylogenies for Cunoniaceae and sections thereof (Table 1). Added genera were placed halfway along the branch leading to their sister genus or clade in the phylogeny. Branch lengths leading to added genera were set such that the tree remained ultrametric. Then, we randomly added species without sequence data into their respective genus.

**Table 1.** Genera added to the molecular phylogeny in this study, because they were present in the distributional dataset, but lacked DNA sequence data to be placed in molecular phylogeny we generated. The position is represented in relation to the most closely related genera following the cited reference.

| Genus | Phylogenetic position | Reference |
| --- | --- | --- |
| *Ackama* | (*Ackama*, *Spiraeopsis*) | [13] |
| *Acrophyllum* | (*Acrophyllum*, *Eucryphia*) | [49] |
| *Acsmithia* | (*Acsmithia*, *Spiraeanthemum*) | [13] |
| *Aistopetalum* | (*Aistopetalum*, *Hooglandia*) *weakly supported | [49] |
| *Bauera* | (*Bauera*, *Ceratopetalum*) | [49] |
| *Opocunonia* | (*Opocunonia* (*Ackama*, *Spiraeopsis*)) | [13] |

### Analyses

To assess tropical or extratropical associations at deeper evolutionary levels within our phylogeny, we reconstructed ancestral states according to our ecological classification for species. These analyses were conducted at the genus-level because the relationships of many species within genera are uncertain, particularly for un-sequenced species, while we have greater confidence in the relationships amongst genera (the majority of nodes in the phylogeny deeper than the genus-level have maximum likelihood bootstrap support values ≥ 70). Genera were classified as Tropical if all the species within a given genus were classified as Tropical (10 genera). Those genera with all their species classified as Extratropical were classified as Extratropical (7 genera). Otherwise, genera were considered Generalist (11 genera). Prior to ancestral state reconstruction, we evaluated support for an equal or unequal transition rate model (for transitions among the three states), with the equal transition rate model best supported. Ancestral states were then estimated using a maximum likelihood with an equal rates model [27]. In order to illustrate the distribution of genera across environmental space, we show the frequency of classified species occurrences across gradients of mean annual temperature (MAT) and mean annual precipitation (MAP).

We estimated taxonomic and phylogenetic indices of diversity for the groups of species classified as “Tropical”, “Extratropical” or “Generalists”. For phylogenetic indices of diversity, we estimated Phylogenetic Diversity (PD, [50]) and Time Integrated Lineage Diversity (TILD, [51]). As the number of lineages monotonically increase towards the present, PD is weighted heavily towards lineage diversity in recent evolutionary time compared to deeper evolutionary time. TILD is estimated from a log-transformed number of lineages at each point in time, which down-weights the influence of recent lineage diversity and gives greater weight to deep-time lineage diversity. The indices therefore can be considered complementary when making comparisons of phylodiversity across groups of species or regions [51]. We also compared values of standardized Phylogenetic Diversity (sPD) to evaluate the phylogenetic structure of the tropical, extratropical and generalist assemblages as a whole. We investigated phylogenetic signal for tropical or extratropical affinity of species using the *D* index [52] for binary traits (with 1,000 permutations in each case to test for significance). Lastly, in order to illustrate variation in lineage diversity over time, we also constructed lineage through time (LTT) plots for a phylogeny comprising all species in tropical regions versus a phylogeny comprising all species occurring in extratropical regions.

In order to evaluate the relationship between species richness, phylogenetic structure and environmental conditions, we arrayed our distributional dataset across an environmental space defined by Mean Annual Temperature (MAT) and Mean Annual Precipitation (MAP). Each species occurrence was assigned to a ‘bin’ coordinated by evenly partitioned vectors of MAT and MAP (60 and 40 bins for each vector, respectively). Thus, we created 2,400 potential bins of species in the environmental space. Many combinations of MAP and MAT do not have any Cunoniaceae species, so not all bins are filled. For each of the bins with species, we estimated species richness and calculated standardized Phylogenetic Diversity (sPD). We efficiently derived the sPD index for so many bins by calculating the first two moments of the null expectation for Phylogenetic Diversity (PD), and then using the moments to calculate a standardized effect size of PD for each bin, which we equate with sPD [53]. Considering the preponderance of *Weinmannia* in the family (133 spp out of 302 spp in our dataset), we estimated these indices on datasets both including and excluding this genus.

We assessed the significance of phylogenetic clustering or overdispersion for species found in each environmental bin by comparing the observed sPD values to the distribution of these values under a null model scenario. We did this because of concerns around artefactual correlations between species richness and sPD values [54]. To develop the null expectation for sPD values, we maintained the total species richness of each bin while randomly assigning species identities by sampling without replacement from the pool of species present in the community matrix. We created 1,000 null assemblages per environmental bin. This null model assumes that all species present in our dataset are equally able to colonize any environmental bin. Bins were considered to be significantly phylogenetically overdispersed or clustered if they occurred in the lowest or highest 2.5% of the distribution of values from the null assemblages (*i. e.*, a two-tailed test with *α* = 0.05).

All analyses were conducted in the R Statistical Environment [36] using functions in the “geiger” [55], “ape” [56], “caper” [57], “Picante” [58], “biogeo” [59] and “PhyloMeasures” [53] packages. A repository with the codes can be found in https://github.com/ricardosegovia/Cunoniaceae.

## Results

Across taxa with enough data to classify, 47 species from 16 genera were classified as Extratropical, 215 species from 21 genera were classified as Tropical and 40 species from five genera were classified as Generalists (Table 2). Eight genera included both Tropical and Extratropical species, while an additional three also included Generalist species. Thus eleven genera in total are classified as generalist. Most of the Generalist species are from the *Weinmannia* genus (33 of 40) and are found in the Andes (Fig. 1), which represent a mosaic of regularly freezing and non-freezing environments. The other genera containing generalist species are *Bauera* (1 sp), *Schizoemeria* (1 sp), *Spiraeanthemum* (2 spp) and *Lamanonia* (3 spp). Tropical species are largely distributed in northern Australia, the Malay Archipelago (plus New Guinea) and Madagascar (Fig. 1). In contrast, Extratropical species are associated with southern Australia, New Zealand, South Africa and southern South America (Fig. 1). Interestingly, *Weinmannia* is absent in Australia and is the only representative of the family in Madagascar (Fig. 1).

**Table 2.**
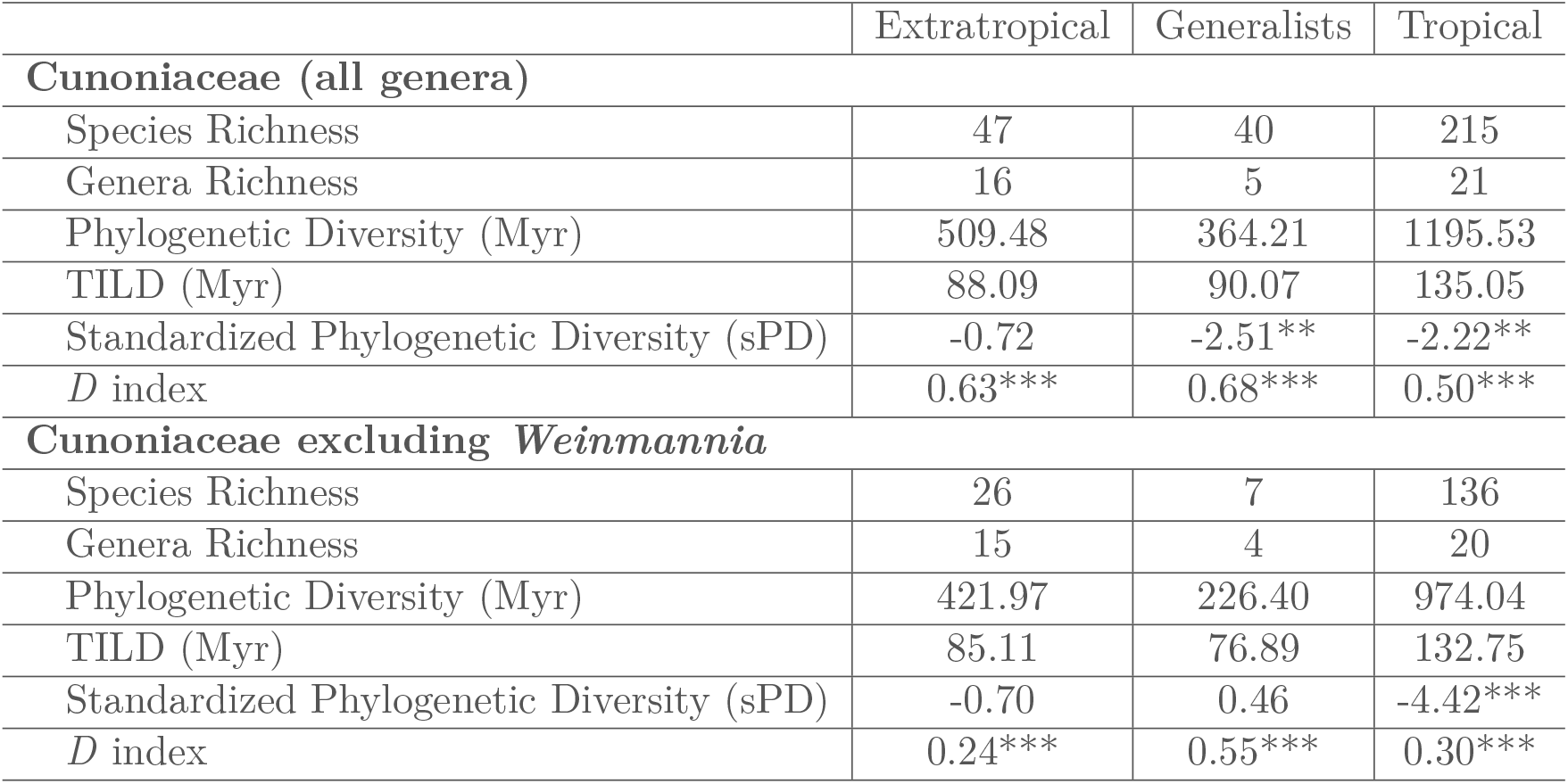
Taxonomic and phylogenetic diversity indices for Tropical, Extratropical and Generalists species assemblages. Some genera contain Tropical and/or Extratropical and/or Generalist species, and are included under more than one column for genus richness totals, as pertinent. The indices are estimated for the whole Cunoniaceae family and for Cunoniaceae excluding *Weinmannia*. For standardized phylogenetic diversity (sPD), asterisks represent the P-value (quantile) of observed PD vs. null communities produced by 1,000 randomizations. For *D*, asterisks represent the probability of estimated *D* resulting under no phylogenetic signal. * if p < 0.1, ** if p < 0.05, *** if p < 0.001.

Ancestral state reconstructions cannot resolve a tropical or extratropical origin for the oldest nodes in the Cunoniaceae family tree. For example, the sister clade to the remainder of the family, composed of *Spiraeanthemum* and *Acsmithia*, has a high association with tropical environments, while the next sister clade of the remainder of the family, composed by *Davidsonia*, *Schizoemeria*, *Ceratopetalum*, *Bauera*, *Platylopus*, and *Anodopetalum*, tends more towards an association with extratropical environments (Fig. 2). In more recently derived clades, ancestral state probabilities show environmental associations that have moved in both tropical-to-extratropical, and extratropical-to-tropical directions (Fig. 2). The high number of genera (11 of 28) with species in both the Tropics and Extratropics further points to the high number of evolutionary shifts between tropical and extratropical environments. Tropical species tend to be distributed in areas with ≥20°C mean annual temperature (MAT), although species from *Spiraeopsis* and *Opocunonia* can occur at lower MAT (Fig. 2). On the other hand, Extratropical species are distributed in environments with lower precipitation, especially the African genera *Cunonia* and *Platylophus* (Fig. 2).

**Figure 2.**
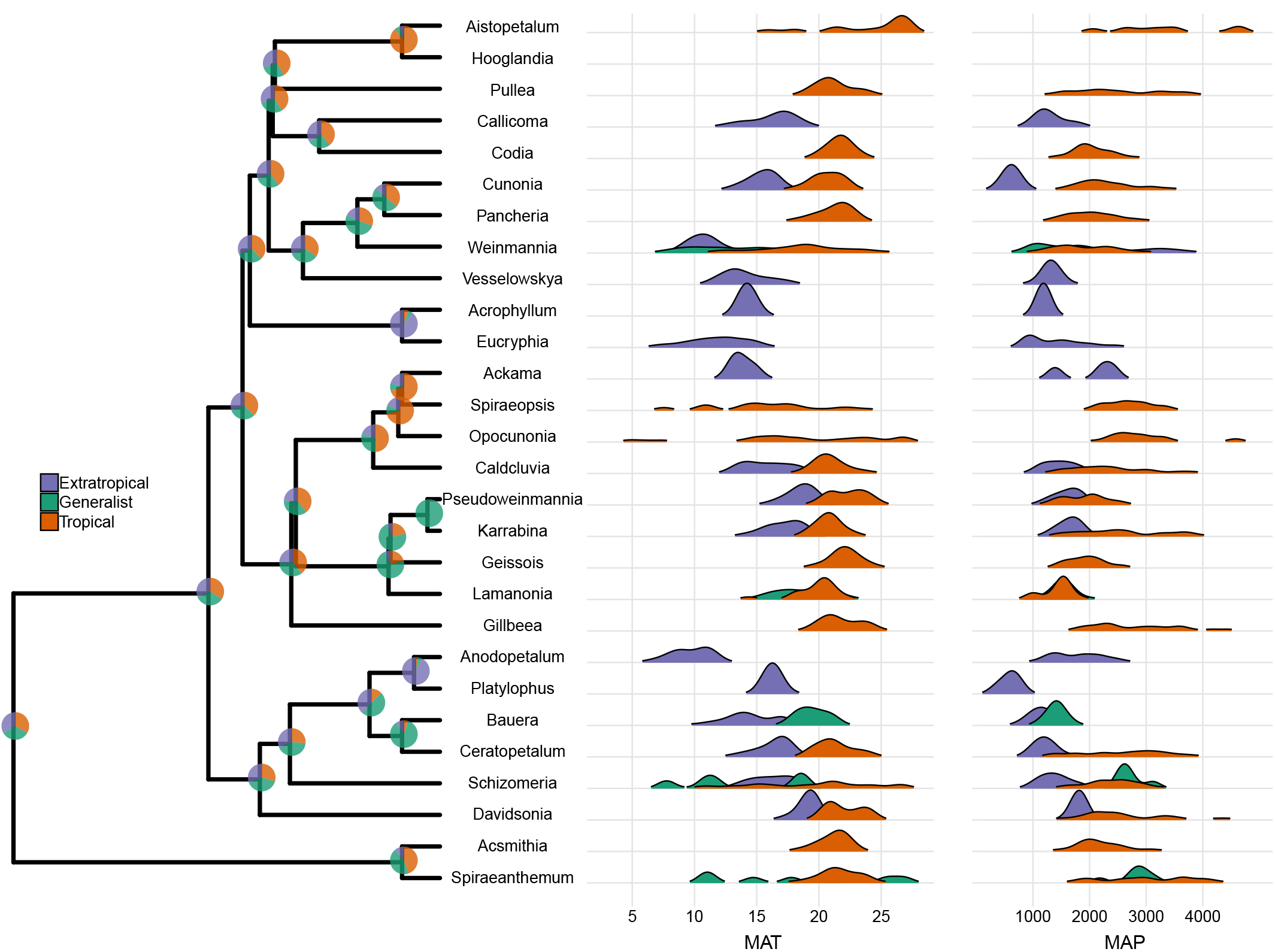
Genus-level phylogeny with ancestral state reconstruction for tropical or extratropical environments. The environmental distributions of Cunoniaceae genera across gradients of Mean Annual Temperature (MAT) and Mean Annual Precipitation (MAP) are also shown. Colours in the right-hand panel illustrate the respective distributions for species within the genus classified as Tropical, Extratropical or Generalist.

The lineage through time (LTT) plot shows that both tropical and extratropical regions contain similar diversity in terms of number of older lineages (~47 Ma) (Fig. 3). The tropical assemblage contains greater lineage diversity for intermediate evolutionary ages (between ~47 and ~12 Myr old). Within the last 12 Myr, the number of lineages in tropical regions has increased much faster than in extratropical regions (note log-scale of y-axis in Fig. 3), which is epitomized by the fact that at present, tropical regions have nearly three times as many species as extratropical regions (245 versus 87 species, when including generalists in both).

**Figure 3.**
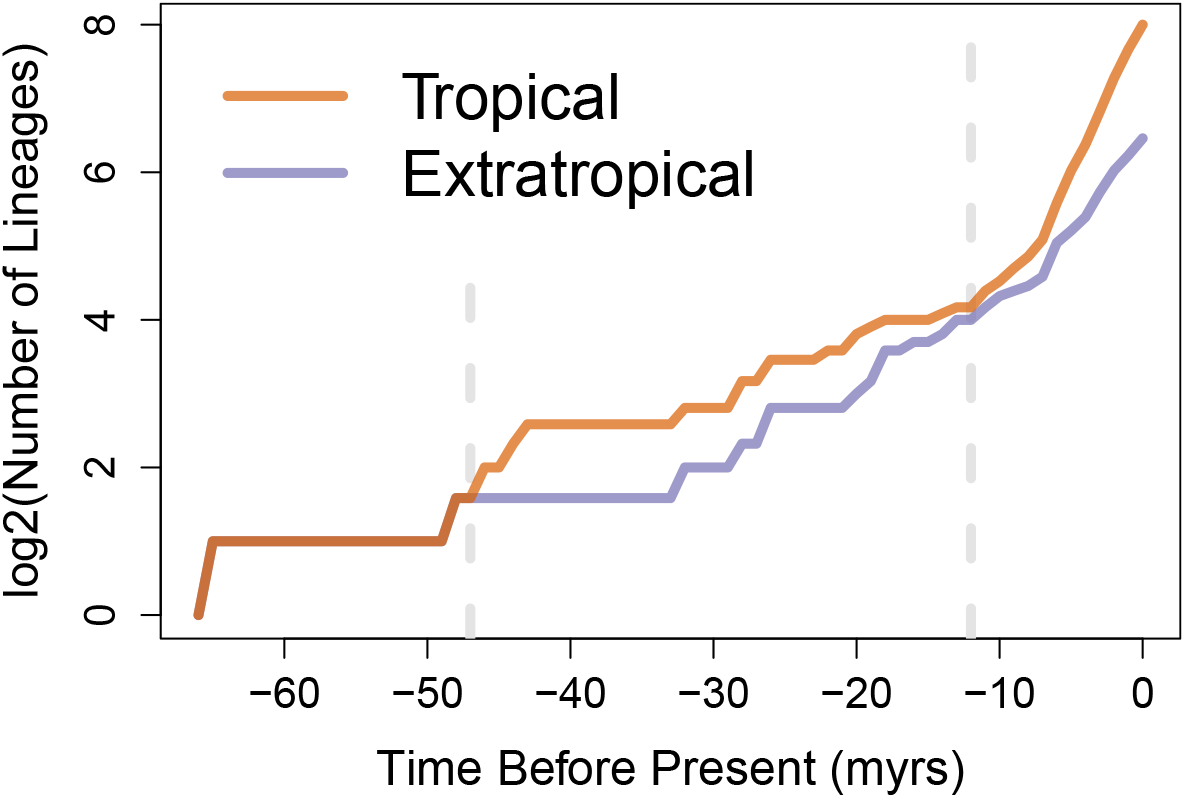
Number of Lineages Through Time (LTT) for Tropical and Extratropical species assemblages. Lineages from Generalist species have been included in both Tropical and Extratropical assemblages. Vertical lines indicate evolutionary depths at which Extratropical and Tropical Assemblages had a similar numbers of lineages.

Overall phylogenetic diversity (PD) is markedly higher for the assemblage of species in tropical environments (non-freezing) than in extratropical environments (freezing; Table 2). The TILD measure of deeper-time lineage diversity is also higher in the tropics, but the difference between tropics and extratropics for that measure is less pronounced. In contrast, sPD is lower, or more negative, for the tropical assemblage than the extratropical assemblage of species (Table 2), indicating greater phylogenetic clustering in the tropics. While the pattern of overall tropical clustering is stronger when excluding *Weinmannia*, the trend of overdispersion in the Generalist group disappears when *Weinmannia* is excluded (Table 2). As such, the generalist distribution of the very speciose *Weinmannia* may be strongly affecting phylodiversity indices and obscuring the patterns of phylogenetic structure observed across the remainder of Cunoniaceae. Complimentary to the above analyses, we find phylogenetic signal, significantly different from random expectations (which would give a *D* value of 1), for both tropical species (*D*= 0.5, *p* < 0.001) and extratropical species (D= 0.64, *p* < 0.001). A lower *D* value in the Tropics reinforces that tropical species are more closely related to each other on average than extratropical species (as values closer to zero indicate greater phylogenetic signal; Table 2).

The distribution of species richness (SR) in environmental bins displays two distinguishable areas of high values (Fig. 4a, d). One area of high species richness is distributed towards relatively high Mean Annual Temperature (MAT) and high Mean Annual Precipitation (Fig 4a, d), while a further area of high richness is distributed towards lower MAT and lower MAP (Fig 4a, d). When excluding *Weinmannia*, only the former peak of species richness (high MAT and MAP) remains (Fig. 4d).

**Figure 4.**
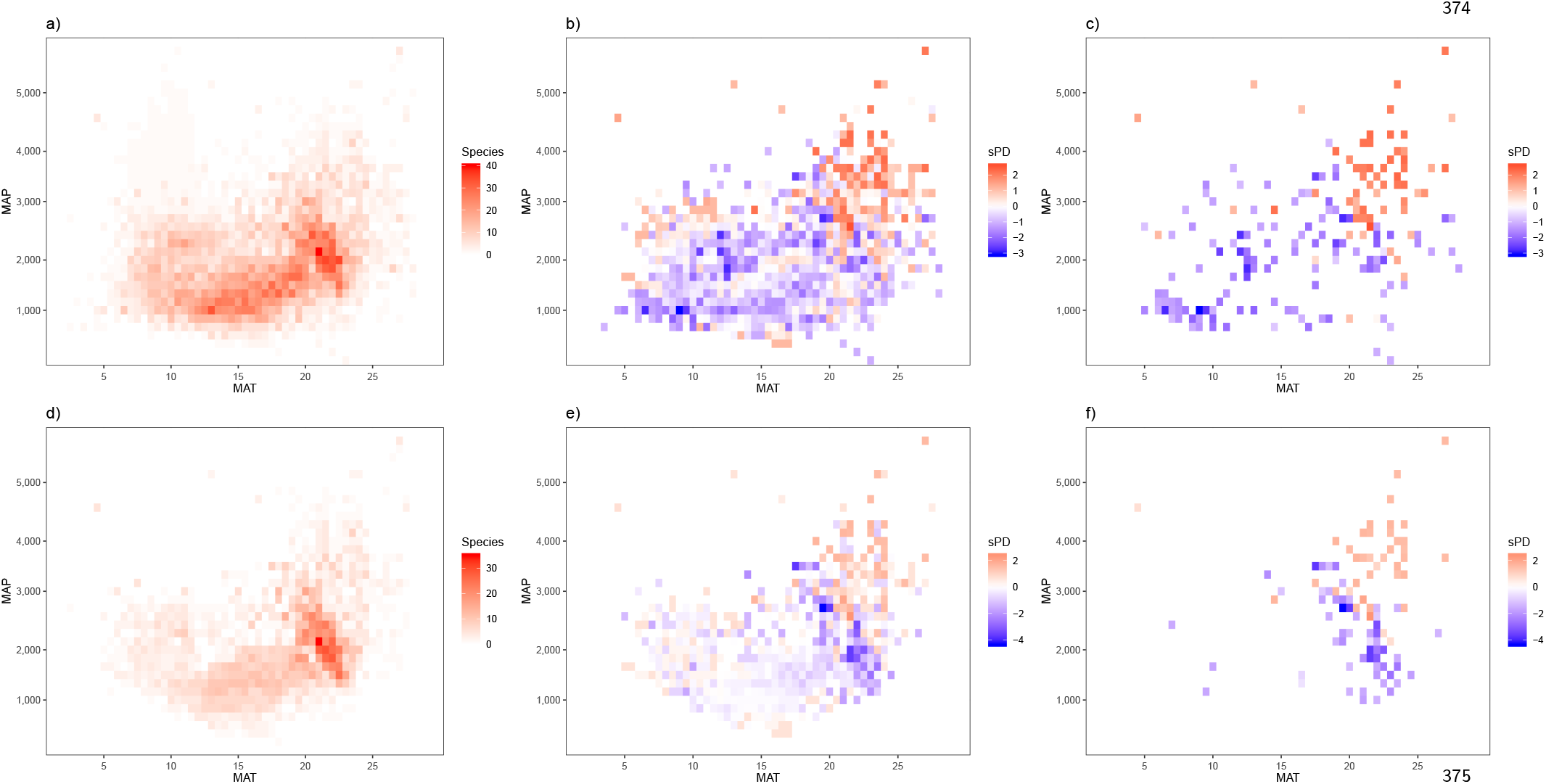
Species Richness (SR) and Standardized Phylogenetic Diversity (sPD) across variation in Mean Annual Temperature (MAT) and Mean Annual Precipitation (MAP). Plots a), b) and c) consider the whole family Cunoniaceae, and d), e) and f) consider the Cunoniaceae family excluding *Weinmannia*. Panels c) and f) show the values of sPD that reflect significant phylogenetic clustering (negative values) or overdispersion (positive values), based on a null model analysis (see main text for details).

A negative correlation (r = −0.313, p<0.0001) was found between SR and standardized Phylogenetic Diversity (sPD) when considering all Cunoniaceae species (Fig. 4a b) and when excluding *Weinmannia* (r = −0.401, p<0.001) (Fig. 4d e). The correlation between SR and sPD suggests that the most speciose assemblages are also the least phylogenetically diverse assemblages. Our null model analysis (Fig. 4c f) allows us to rule out a potentially artefactual relationship between SR and sPD, as cautioned by [54], at least for the extremes of species richness values. When considering the whole family (Fig 4a c), we see that the most phylogenetically overdispersed assemblages can be found within the hottest and wettest environments, while a number of phylogenetically clustered assemblages are found towards cooler and drier environments (in both tropical and extratropical portions of climatic space). When *Weinmannia* is excluded from analyses (Fig. 4e f), those environmental bins that show significant phylogenetic clustering are in relatively warm and wet (*i. e.* tropical) environments compared to the environmental range of the genus as a whole (Fig. 2). Meanwhile, there are few environmental bins that show significant phylogenetic overdispersion once *Weinmannia* is excluded from analyses. *Weinmannia* seems to constitute an independent pattern of diversification from the rest of the family with respect to environmental gradients and to have a strong influence on overall results (contrast Fig. 4a c versus d f).

## Discussion

The angiosperm family Cunoniaceae has had strong associations with both tropical and extratropical environments over its evolutionary history (Fig. 2), yet tropical environments currently hold many more species. Remarkably, this difference in diversity seems to have arisen within the last 12 Myr (Fig. 3), suggesting higher recent diversification in the tropics relative to the extratropics. We also find significantly less phylogenetic diversity in tropical regions than expected given their species richness (*i. e.* significant phylogenetic clustering; Fig. 4, Table 2), suggesting that recent radiations have greatly increased the number of species in the tropics without substantially increasing phylogenetic diversity.

The tropical radiations that comprise the majority of Cunoniaceae species may have arisen from extatropical ancestors. If so, this would belie the general angiosperm pattern of species-poor extratropical lineages descending from tropical ancestors [9]. Rather, it would suggest that that extratropics have been an important provider of diversity in tropical regions. However, the frequent switching of Cunoniaceae lineages between the extratropics and tropics limits our ability to reconstruct ancestral environmental preferences of lineages (Fig. 2), at least from distribution data for extant taxa. The fossil record indicates that the family Cunoniaceae first appeared during the Late Cretaceous in Antarctica (~70 Ma, [60]. During this time, Antarctica supported a highly diverse vegetation, dominated by *Podocarpites* and *Nothofagus* [61], which is similar in taxonomic composition to today’s Valdivian rainforest in Chile ([48]; and references therein). Although there are no climatic models which suggest whether Antarctica experienced freezing or non-freezing conditions 70 Ma, the compositional similarity with extant Valdivian forests, which do experience freezing, suggests that the ancestor of present-day Cunoniaceae lineages originated in an environment with freezing temperatures.

Based on distribution data from extant taxa, a tropical or extratropical origin for the most speciose genus in the family, *Weinmannia* is also not clear (Fig. 2). A morphological and molecular cladistic analysis shows that *Weinmannia* section Leiospermum, from extratropical New Zealand, tropical New Caledonia, and other islands from the South Pacific, is the sister taxon to the remainder of the genus [62]. These tropical and extratropical affiliations in the basal section of *Weinmannia* could indicate an origin of the clade associated with the interface between freezing and non-freezing environments. Meanwhile, in section Weinmannia, the most recently derived section of the genus, species from the tropical Mascarene islands are the sister clade of the American species [62]. The American species of section Weinmannia represent an impressive radiation that coincides with the Andean uplift (~60 species in the Andes from Argentina to Mexico). Following our ecological classification of species, the section Weinmannia is represented by only one extratropical species (in southern South America), yet those species distributed across the “Tropical Andes” are not inhabiting truly tropical environments (Fig. 1, Table 2). Rather, they occur at the interface between freezing and non-freezing temperatures in mid and high elevations, which is why so many of them are classified as generalists. Meanwhile, the fossil record shows a potential ancestor of section Weinmannia in the extratropical Tasmania [63]. Therefore, an extratropical origin is distinctly possible for the hyperdiverse genus *Weinmannia* as a whole and the recently radiated section Weinmannia.

Patterns of trait variation across species in different environments could also potentially give insights into the biogeographic history of Cunoniaceae. For example, across angiosperms as a whole, a pattern of decreasing hydraulic conduit diameter with increasing latitude is thought to be an adaptive consequence of lineages moving into freezing environments [5, 64]. Evidence for this pattern in *Weinmannia* [65] could be taken to suggest a tropical origin for this genus, with a later dispersal into freezing environments. However, phylogenetic evidence suggests an extratropical origin for *Weinmannia* is as probable as a tropical origin (Fig. 2). We therefore suggest that varying conduit diameter among species may just as likely represent either a plastic response to the presence or absence of freezing conditions or an adaptive trend of increasing conduit diameter when lineages move into tropical environments from extratropical ones. The latter pattern in vessel diameter is evident in *Nothofagus*, a genus whose extratropical origin is more certain. *Nothofagus* species inhabiting tropical environments are only found in New Guinea and have larger vessel diameters than the remaining species of the genus, which inhabit extratropical environments [66]. These patterns in Cunoniaceae and *Nothofagus* suggest that extratropical biota can potentially change their hydraulic architecture when dispersing into tropical, non-freezing environments.

Beyond the origin of Cunoniaceae and its most species-rich genus, the diversification history of the family shows two time periods during which tropical lineages became more numerous than extratropical lineages, beginning ~47 and ~12 Ma (Fig. 3). The start of the first time period is synchronous with the early Eocene Climatic Optimum [67], which is the warmest period of the Cenozoic and may have spurred lineage diversification in the tropics. This warm period ended ~30 Myr before present, corresponding with the inception of Antarctic freezing around the Eocene–Oligocene boundary [67]. This freezing is linked with the extinction of the Antarctic flora, including Cunoniaceae present there [68, 69]. The second time period during which tropical lineages gained predominance started in the late Miocene (after 12 Myr). During this later time period, global temperatures continued to cool, culminating in the Pleistocene glaciations. At this time the southern extratropical regions of the world would have been subject to important environmental changes, including increased seasonality and dryness in southern mid-latitudes, which may have caused a large number of extinctions [70]. Such extinctions could explain the discrepancy in diversification rate between the tropics and extratropics over the last 12 Myr. It is important to keep in mind however, that these differences in lineage diversity over the evolutionary history of Cunoniaceae are inferred from phylogenetic data on extant lineages. The actual lineage diversity of Cunoniaceae in extratropical or tropical regions of the world during past geological epochs may well have been higher, with some lineages not surviving until the present day.

The negative relationship between species richness (SR) and standardized phylogenetic diversity (sPD) observed across environmental bins (Fig. 4) suggests that the environmental peak of SR in the Cunoniaceae family is explained by radiation of a subset of lineages into new environments. Similar reasoning has been applied to other systems. On one hand, phylogenetic clustering (*i. e.* low sPD) in Floridan plant communities has been interpreted as a consequence of habitat filtering for a subset of lineages containing specific, phylogenetically conserved ecological traits [71]. On the other hand, evidence for phylogenetic clustering in the western Cape flora of South Africa has been discussed as a function of multiple, rapid radiations [72, 73]. Indeed, differences in the mean number of species per genus for genera inhabiting the tropics versus the extratropics in Cunoniaceae reinforce the hypothesis of restricted radiation for a largely tropical subset of lineages. While the tropics house 215 species from 21 genera (10.2 spp per genus on average), the extratropics house 47 species from 16 genera (2.9 spp per genus on average) (Table 2). In other words, the relatively few species inhabiting the extratropics come from a roughly similar number of genera as the many species in the tropics.

Higher species richness in the tropics can arise from multiple, non-mutually exclusive mechanisms, such as area effects, time for speciation, stronger biotic interactions or increased mutation rates (reviewed by [74]). In the case of Cunoniaceae, area effects do not seem a feasible explanation because its truly tropical distribution is restricted to Australasia and Madagascar, a much narrower distribution than the combined generalist and extratropical distribution of the family (Fig. 1). An explanation based on time for speciation also seems unlikely, because this would imply a later dispersion of Cunoniaceae into freezing environments and, as we have shown here, an extratropical origin for the family seems as likely (Fig. 2), or perhaps more likely considering the fossil record, than a tropical origin. Furthermore, tropical species are more phylogenetically clustered than extratropical species, which suggests that a tropical distribution may be novel for the family compared to an extratropical distribution. Instead, high tropical species richness in Cunoniaceae may be due to high recent diversification rates in tropical Cunoniaceae lineages, which could in turn be due to a multiplicity of processes including stronger biotic interactions within tropical lineages (*e. g.* [75]) or increased mutation rates linked to higher temperature [76]. Regardless, future studies are clearly needed to address the mechanisms behind the radiation of Cunoniaceae into non-freezing environments.

The prominent radiation of *Weinmannia* in the Andean mountains is not easily reconciled with environmental factors generally considered to increase speciation rate. An historical perspective may work better. The Andes are not older than the South American extratropics, yet they are much more diverse (~60 vs. 1 spp of *Weinmannia*) [77], making time for speciation explanations unlikely. Higher mutation rates due to increased temperatures and/or solar radiation are also not feasible, because *Weinmannia* prefers relatively colder and cloudy environments in the Andes [78]. An explanation for the radiation based on competition does not represent a strong hypothesis either, since *Weinmannia* is a pioneer genus and a weak competitor [79]. In contrast, the high taxonomic diversity in low latitudes of the Andes may be associated mainly with topographic dynamism during the relatively recent Andean uplift [80, 81].

Our findings present an explanation for Southern Hemisphere biogeographic trends which goes beyond the traditional dispersal-vicariance dichotomy by explicitly focusing on the environmental context under which lineages evolved. The family Cunoniaceae is of clear importance, in terms of diversity and abundance, in both tropical and extratropical regions. As such it serves as a model family for understanding the nature of shifts between tropical and extratropical environments, and their consequence for diversification. In contrast to the general pattern in angiosperm biogeography [9, 5], we find that Cunoniaceae lineages have made frequent shifts between tropical and extratropical environments. Further, while the general narrative for biogeography in angiosperms is one of tropical-origin lineages spreading into the extratropics [8], the New Caledonian and Indonesian radiation of Cunoniaceae and the radiation of *Weinmannia* in the tropical Andes, may represent exceptions to this rule. These radiations seem to have extratropical origins or at least to come from lineages that have continually existed at the extratropical-tropical interface. Therefore, the dispersal of Cunoniaceae lineages into tropical biomes represents a striking case of non-tropical biota contributing taxonomic and evolutionary diversity to tropical biodiversity hotspots.

## Acknowledgements

CONICYT PIA APOYO CCTE AFB170008. R.A.S. is supported by a Newton International Fellowship from The Royal Society and by Conicyt PFCHA/Postdoctorado Becas Chile/2017 N° 3140189. A.R.G. is supported by a NERC studentship (NE/L002558/1). K.G.D. was supported by a Leverhulme International Academic Fellowship during the time this research was completed.

